# Female-female aggression in a sexual/unisexual species complex over resources

**DOI:** 10.1101/057018

**Authors:** Amber Makowicz, Tana Moore, Ingo Schlupp

**Affiliations:** Department of Biology, University of Oklahoma, 730 5 Van Vleet Oval, Norman, OK 73019, USA; Department of Biology, Lehrstuhl für Zoologie und Evolutionsbiologie, University Konstanz, Universitätsstraβe 10, 78457 Konstanz, Germany

**Keywords:** female-female aggression, food resource, kin selection, *Poecilia formosa*, *Poecilia latipinna*, sexual/unisexual mating system

## Abstract

Kin selection theory predicts that closely related individuals should be more altruistic and less antagonistic towards one another. In addition, it would predict that the higher the relatedness value (R) between individuals, such as in clonal organisms, the more likely kin selection is to evolve. One benefit of kin selection is a reduction in aggression towards kininvarious social contexts, such as foraging. In the gynogenetic Amazon molly, *Poecilia formosa,* females have been shown to differentiate between clone types, preferring to associate with clonal sisters to non-sisters, and regulate their aggressive behaviors accordingly. We ask if Amazon mollies in resource-limited environments: 1) still maintain the ability to regulate aggressive behaviors according to relatedness (heterospecific females, clonal sisters or non-sisters), and 2) how their aggressive behaviors change relative to a female’s social partner? Using a repeated-measures design, we found that focal females regulated their aggressive behaviors depending on partner type (i.e., a heterospecific female, clonal sister, or non-sister). Heterospecific females and the non-sister clones spent more time behaving aggressively towards the focal females, and these females also received significantly more bites from heterospecific females. Interestingly, the clonal sisters, when compared to the other two female types, performed significantly more tail beats towards focal females.We are able to confirm that females do show more aggression towards heterospecific females and non-sister clones in a food-limited environment, and that their aggression scales with relatedness (R).

**Lay Summary:** Kin recognition allows individuals to adjust costly behaviors, such as aggression, according to the degree of genetic relatedness. We show that in a food-limited environment, a clonal species of fish, the Amazon molly, females regulate aggressive behaviors depending on how closely related they are to the recipient female, behaving more aggressively to both heterospecific females and non-sister clones. The ability to regulate costly behaviors under variable socialconditions is advantageous, especially when resources are limited.

## Introduction

### Kin recognition

The theory of kin selection predicts that closely related individuals benefit from the reproductive fitness of one another; therefore, closely related individuals should be more altruistic and less antagonistic (Hamilton 1963). The concept of inclusive fitness was introduced by Haldane and later formalized by W. D. Hamilton (Hamilton’s rule; Hamilton 1963). Hamilton’s rule was used as a powerful tool to explain seemingly altruistic behaviors, especially in eusocial hymenoptera (although kin selection is by no means restricted to hymenopterans; Waldman 1988; Pfenning et al. 1993). Although there has been some debate in recent literature (Nowak et al. 2010; Abbot et al. 2011; Boomsma et al. 2011; Ferriere and Michod 2011; Herre and Wcislo 2011; Strassmann et al. 2011), this theory still remains prominent when investigating the evolution of particular behaviors like, cooperation, reduced aggression, and more.

In sexual species, because of meiosis, direct descendants share only 50% of their genes with either the mother or father. As such, the majority of research has investigated how distant individuals must be for kin selection to occur (Waldman 1988; Pfenning et al. 1993). Clonal animals present an interesting paradox: although clonality leads to R-values of up to 1 (as in identical twins), the differences between distinct clonal lineages are often very minute. This raises interesting questions concerning the power of kin selection in clonal organisms: *how small* can the genetic differences be between individuals while still allowingforadaptive kin recognition to evolve?

Theory would predict that the higher the relatedness value (R) between individuals, the more likely kin selection is to evolve. However, kin recognition in clonal species (or clonal selection) has produced mixed results. In parthenogenetic ants (*Cerapachys biroi*), colonies are composed of identical sisters that can distinguish and select against non-sister clones and even different colonies of the same clone and therefore, are able to maintain a colony that lacks internal conflict (Kronauer et al. 2013). In polyembryonic parasitoid wasps (*Phytoseiulus perimilis*), hundreds of offspring can be produced from a single fertilized egg, and soldiers are able to identify and attack non-sister clones, thus reducing competition for their clonal sisters within a parasitized caterpillar (Segoli et al. 2009). However, cyclical parthenogenetic (individuals that reproduce both sexually and clonally depending on environmental cues) water fleas (*Daphnia pulex*), and social aphids (*Tamalia coweni*), do not seem to recognize identical sister clones (Winsor and Innes 2002; Miller III 1998). Finally, polyembryonic armadillos (*Dasypus novemcinctus*) have 4 identical offspring in each litter, and these young do not recognize identical siblings either as juveniles or adults (Loughry and McDonough 2013).

There are several parameters that must emerge prior to the evolution of kin selection, and it is hypothesized that lacking in one or more parameter is why we see such variation within clonal species (Loughry and McDonough 2013). Kin selection is likely to evolve when: 1) individuals have the ability to discriminate between clonal sisters and non-sisters independently of context and familiarity, 2) the benefits favoring clonal sisters override the costs to self-fitness, 3) multiple clonal lineages have overlapping social environments in both time and space, 4) dispersal of the clones is not too great a distance to prevent overlap, and 5) aggregations of individuals are large in size (Waldman 1988; Loughry and McDonough 2013). Thus far for kin selection in clonal animals, the negative results in some clonal invertebrates fail one or more of these assumptions. For instance, *Daphnia* have wide dispersal ranges with the likelihood of encountering an identical sister being slim (Winsor and Innes 2002).The social aphid, *Tamalia coweni,* live in galls that typically only contain clonal sisters, and the frequency of encountering non-sister clones is rare (Miller III 1998); whereas the non-social pea aphids (*Acyrthosiphon pisum*) encounter both clonal sisters and non-sisters and do show kin recognition (Muratori et al. 2014). Even inthe polyembryonic armadillo, they disperse far and they do not form social aggregations as adults (Loughry and McDonough 2013). Furthermore, to date, there are no studies on true clonal vertebrates. In the present study, we use the gynogenetic Amazon molly, *Poecilia formosa*, which meets several of the above mentioned assumptions: multiple clonal lineages in an overlapping social environment, with limited dispersal, and large aggregations.

### Sexual/unisexual mating system

Amazon mollies are gynogenetic hybrids that live in syntopy with either of their parental species, the sailfin molly (*Poecilia latipinna*) and the Atlantic molly (*Poecilia mexicana*) (Amazon hybrid origin c.a. 120,000 generations ago; Schartl et al. 1995; Stöck et al. 2010). Gynogenetic females require sperm from a sexual host, commonly either parental species theAtlantic or sailfin molly males, to initiate embryogenesis (Hubbs and Hubbs 1932); however, the male’s DNA is not incorporated into the genome of the offspring, resulting in identical sister clones (Schlupp 2005). The genome of the species is very homogenous and all known genetic variation is due either to mutation, gene conversion,or introgression of paternal DNA (Schlupp and Riesch 2011). Overall, relatedness within the species is very high, even among different clonal lineages.

Amazon mollies occur with their sexual hosts in large, open shoals that fluctuate in the ratio of each species during the breeding season (Heubel 2004). As such, they tend to compete with the sexual females of the parental species for many resources, and overlap in many aspects of their ecological niche, life history parameters, and behaviors (Heubel 2004; Tobler et al. 2005; Hubbs and Schlupp 2008; Riesch et al. 2008; Fischer and Schlupp 2010; Schlupp et al. 2010; Tobler and Schlupp 2010; Scharnweber et al. 2011a; Scharnweber et al. 2011b; Scharnweber et al. 2011c; Alberici da Barbiano et al. 2014). It has been proposed by the ‘behavioral regulation hypothesis’ that the stability between host and sexual parasite is maintained by adaptive male mate choice and is thus maintaining this species complex (Schlupp and Riesch 2011). Males have a preference for large, conspecific females, but will mate with Amazon mollies in order to increase the likelihood of conspecific females copying the mate choice of the Amazons (Schlupp et al. 1994). Furthermore, sexual females are significantly more likely to have sperm in their genital tract and to be pregnant than the unisexual females (Riesch et al. 2012). It has been proposed that counter-adaptations,such as circumventing male choice via aggression towards the preferred sexual females, may allow Amazon females to thwart male mate choice (Schlupp et al. 1991; Heubel and Plath 2008; Makowicz and Schlupp 2015). For instance, Amazon mollies were found to chase sexual females, both sailfin and Atlantic mollies, away from the males (Schlupp et al. 1991).

Further investigation into the aggressive behaviors of Amazon mollies show that aggression in Amazons is a costly behavior. Amazon aggressor females had lower body condition than recipient conspecifics, although this was not the case in sailfin molly females (Makowicz and Schlupp 2015). Amazons tend to behave more aggressively towards their partner, and increase their aggressive behaviors over time more so than sailfin females. Small Amazon molly females were more aggressive towards larger females partners than small sailfin females (Makowicz and Schlupp 2015). In addition, Amazon females tend to form dominance hierarchies from social interactions occurring early in life, with dominant females performing more bites towards subordinate females (Laskowski et al. 2016). These results suggest that there must be some advantage to the high cost of aggressive behaviors found in Amazon mollies.

### Clonal recognition in Amazons

Recent work has shown that Amazon mollies are able to recognize and prefer clonal sisters to non-sisters (Makowicz et al. 2016). The existence of kin recognition in this species is interesting in and of itself, however, the study also showed that Amazon females could discriminate between clonal types using visual only and chemical only cues. Indeed, there was no difference between unimodal (visual or chemical only) or bimodal (the combination of visual and chemical) cues. This was confirmed in a field based experiment, where in highly turbid environments, Amazon mollies depended on chemical cues more readily than in the laboratory, where visual cues were more predominant (Makowicz et al. 2016). Makowicz et al. (2016) predicted that one of the adaptive values of this behavior was to regulate the aggressive behaviors found in this species. Amazons would behave more aggressively towards non-sister clones than clonal sisters. In this current study, we would like to understand the different conditions in the social environment in which females would regulate their aggressive behaviors, particularly if aggression is still regulated when females are competing for a limited food resource.

There are several studies demonstrating that aggression between individuals increases as the availability of food resources decreases (Lim et al. 2014; Grant et al. 2002), or as the time spent with individual conspecifics decreases (i.e., more aggressive towards unfamiliar as compared to familiar individuals; Utne-Palm and Hart 2000). Similar results have been found when relatedness is considered, either on an individual level or in groups (Brown and Brown 1996; Griffiths and Armstrong 2002; Olesen and Jarvi 1997). In Arctic charr, *Salvelinus alpinus,* the frequency of agonistic acts was significantly higher in mixed groups as compared to pure sibling groups after feeding ( Olesen and Jarvi 1997). Atlantic salmon, *Salmo salar,* and Rainbow trout, *Oncorhynchus myki,* both exhibita greater number of aggressive interactions with non-kin, and thus kin-biased foraging behavior facilitates decreased levels of aggression in both species (Brown and Brown 1996). Finally, in Japanese macaques (*Macaca fuscata*) aggression exhibited by dominant females increased significantly with decreasing degree of relatedness in the overall average number of aggressive acts, and with increased distance from the food resource (Belisle and Chapais 2001). Although, the intensity level of aggression by dominant females was not influenced by relatedness (Belisle and Chapais 2001). However, this is not always the case, for instance, in European earwigs, *Forficula auricularia,* neither relatedness nor food deprivation affectedthe durationofaggression (Weib et al. 2014). Given the relationship found between aggression, food availability, and relatedness, we ask if Amazon mollies are more aggressive towards heterospecific females, clonal sister or non-sisters in low food resource environments? We predict that if females are in a food resource limited environment, aggression would still be regulated and that they would show more aggression (via bites and tailbeats) towards heterospecifics than conspecifics, and towards non-sisters than clonal sisters.

## Methods

### Fish maintenance

The focal female population of Amazon mollies used here was derived from a single female that was collected in 2011 from Comal Springs (29°42’46.86” N; 98°08’8.57” W) in New Braunfels, Texas. Microsatellites were used to confirm clonality of the population (i.e., that all individuals within the population were identical to each other, genetic identity=1.0; Makowicz et al. 2016). Stimulus female sailfin mollies were descendents from females collected in 2011 fromComal Springs. Stimulus non-sister clones (genetic identity=0.997) were derived from a single female originating from Rio Purificacion, Barretal, Mexico (24°4’42.85” N; 99°7’21.76” W; collected 2009). The relatedness value (R) between the focal females/clonal sisters (Comal Spring)and the non-sister clones (Barretal) is-0.072, suggesting that these females are as genetically distinct from each other as sexual non-kin individuals. All three populations were maintained in 1000L stock tanks at the Aquatic Research Facility on the University of Oklahoma campus in Norman, Oklahoma.

Prior to the experiment, individuals were transferred into the laboratory and maintained separately in several 50×30×25 cm aquaria (length×width×height). Two weeks before the tests, individuals from the focal population were randomly selected and transferred to smaller (3.8 L) isolated tanks where visual contact with other fish was prevented. Fish were labeled as either focal female or sister clone. Stimulus females were also transferred to isolated tanks and labeled by their population. Standard length (mm) was measured for each individual, both focal and stimulus females, in order to form *a priori* size-matched pairs. All females were maintained at a temperature of 27°C, under a 12:12-hour light:dark cycle, and fed daily *ad libitum*with commercial flake food (Tetramin flakes) and bloodworms. All fish were non-virgins, however, all females were in comparable reproductive states (not pregnant) for the duration of this experiment.

### Experimental design

A foraging experiment allowed an Amazon female to forage with 3 partner types: 1) a heterospecific female, *P. latipinna*; 2) a conspecific female, clonal sister; and 3) a conspecific female, non-sister clone. Focal females were food deprived for 24 hours prior to the experiment, and the partner females were fed daily, including the morning of the experiment. The experimental tank contained clean water and a food tablet that was placed in the center, front of the experimental tank. Stimulus and focal females were placed in clear Plexiglas cylinders in the center of the tank. After a 5-minute acclimation period, both females were released fromthe Plexiglas cylinders and allowed to forage for 5 minutes. Aggressive behaviors (bites, tail beats, and total time spent being aggressive, including the time spent chasing the other female) were recorded. After the treatment concluded, the focal females were placed back into their isolation tank and fed. The next day the focal females were given a recovery day andfednormally. Females were then retested 48hrs after the first treatment with a different partner until all 3 treatments were complete (N=40).

### Statistical Analysis

A repeated-measures GLM was used to analyze the differences in the aggressive behaviors and reception of the behaviors between the three treatments. Within subject variables included: “Treatment” (Heterospecific, Clonal Sister, Non-Sister), “Behavior” (Time, Bites,Tail beats), and “Reception” (Given, Received). The model originally included the standard lengths of the focal females, heterospecific females, clonal sisters, and non-sisters as covariates; however, all stimulus female lengths had no significant effect and were removed from the model. We retained the focal female standard length (SL) in the second model, as this was the only covariate that had a significant effect. All statistics were performed in SPSS (ver. 17).

## Results

Standard length and body weight did not differ between the focal females, heterospecific partners, clonal sisters, and the non-sisters (Table 1).The type of partner female (i.e.,a heterospecific female, clonal sister, and non-sister clone) had a significant influence on the aggressive behaviors of the focal females (Treatment: F_35_=3.667, *p*=0.036; Table 2). The partner females also significantly influenced whether or not females gave or received more aggressive behaviors (Treatment*Reception: F_35_=4.867, *p*=0.014) and this influenced what specific behaviors were more pronounced (Behavior*Reception: F_35_=6.080, *p*=0.005; Figure 1–3). There was no significant difference in the amount of time focal females behaved aggressively, although they tended to behave less aggressively towards clonal sisters (Figure 3, Table 3). Focal females also tended to perform more tail beats towards the heterospecific females and non-sister clones and performed significantly more bites towards the non-sister clones (Figure 1). Although partner female standard length had no influence on the data, the focal female standard length tended to significantly influence aggressive behaviors, with larger females behaving more aggressively (Treatment*FocalSL: F_35_=3.141, *p*=0.056; Treatment*Reception*FocalSL: F_35_=4.631, *p*=0.016; Behavior*Reception*FocalSL: F_35_=5.597, *p*=0.008; Table 2). Females typically received more aggressive behaviors from the heterospecific female and the non-sister clone (Figure 3, Table 3) in the total time and they received significantly more bites from the heterospecific females. Intriguingly,focal females received more tail beats from the clonal sisters when compared to the other two female types (Figure 2).

**Table 1:** The average±SD body size of the focal Amazons, the heterospecific females, clonal sister and non-sister clones. Standard length (SL; mm) was measured to maintain a constant measurement of size for pairing of females. Body weight (WT; g) was measured to ensure that females of similar body condition were used throughout the experiment.

| **Females** | **SL** | **WT** |
| --- | --- | --- |
| Focal | $39 \pm 5.375$ | $1.506 \pm 0.779$ |
| Heterospecific | $38.4 \pm 5.878$ | $1.571 \pm 0.799$ |
| Clonal Sister | $39.38 \pm 4.970$ | $1.544 \pm 0.777$ |
| Non-Sister Clone | $38.95 \pm 4.971$ | $1.512 \pm 0.754$ |

**Table 2:** Results from the repeated-measures GLM indicating all the interaction terms. Bold terms indicate a significant p-value.

| Effect | F | DF | *P*-value |
| --- | --- | --- | --- |
| **Treatment** | **3.667** | **35** | **0.036** |
| Treatment*Focal SL | 3.3141 | 35 | 0.056 |
| Behavior | 1.277 | 35 | 0.292 |
| Behavior*Focal SL | 0.966 | 35 | 0.391 |
| Reception | 0 | 36 | 0.997 |
| Reception*Focal SL | 0.23 | 36 | 0.88 |
| Treatment*Behavior | 0.843 | 33 | 0.508 |
| Treatment*Behavior*Focal SL | 0.653 | 33 | 0.629 |
| **Treatment\*Reception** | **4.867** | **35** | **0.014** |
| **Treatment\*Reception\*Focal SL** | **4.631** | **35** | **0.016** |
| **Behavior\*Reception** | **6.08** | **35** | **0.005** |
| **Behavior\*Reception\*Focal SL** | **5.597** | **35** | **0.008** |
| Treatment*Behavior*Reception | 1.857 | 33 | 0.142 |
| Treatment*Behavior*Reception*Focal SL | 1.542 | 33 | 0.213 |

**Fig. 1.**
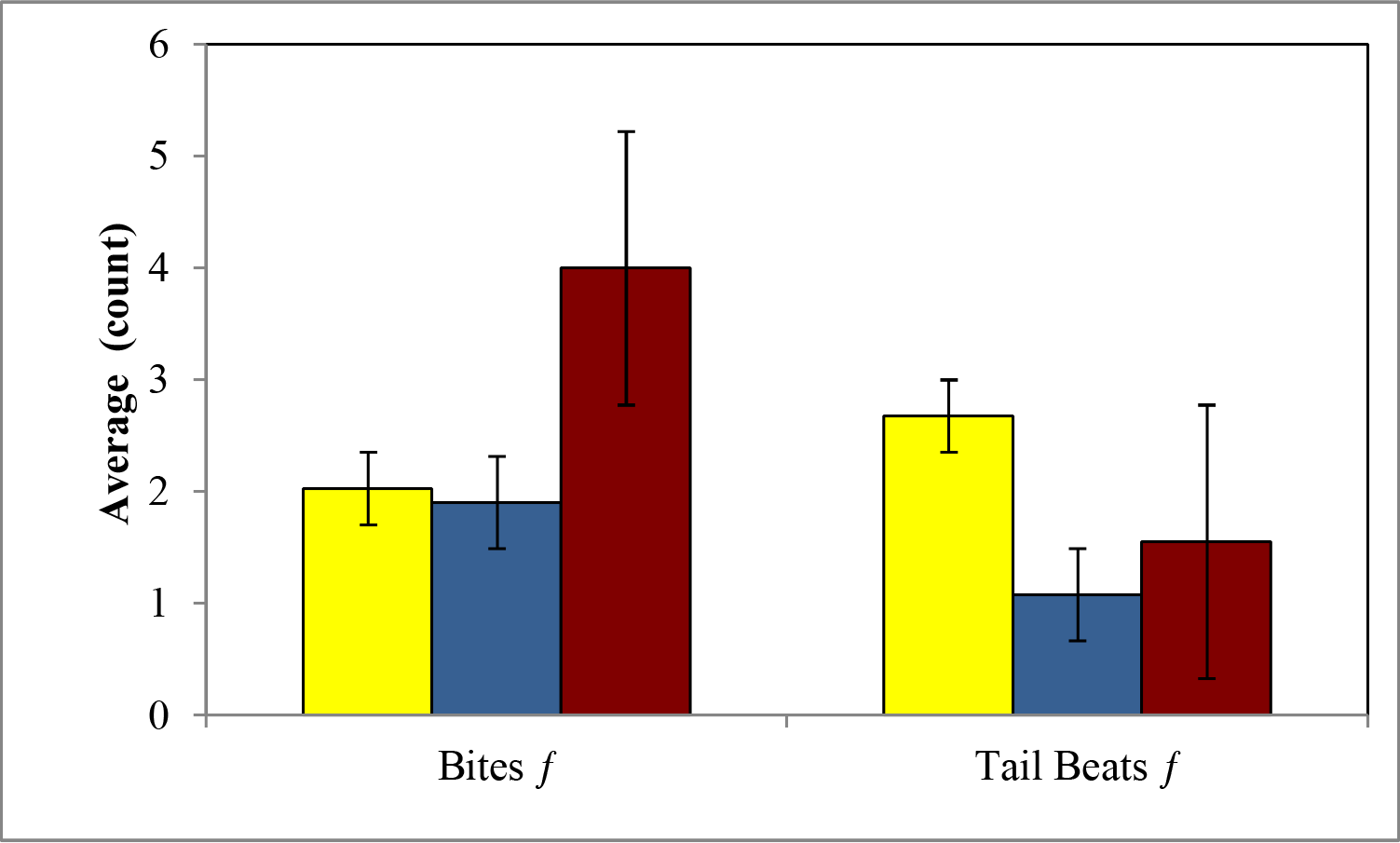
Average±SE amount of aggression in bites and tail beatsgiven towards partner females: heterospecific females (yellow), clonal sisters (blue), and non-sister clones (red). Focal females gave more bites tonon-sister clones when compared to heterospecific females or clonal sisters. Additionally, females gave more tail beats to heterospecific females when compared to either clone type.

**Fig. 2.**
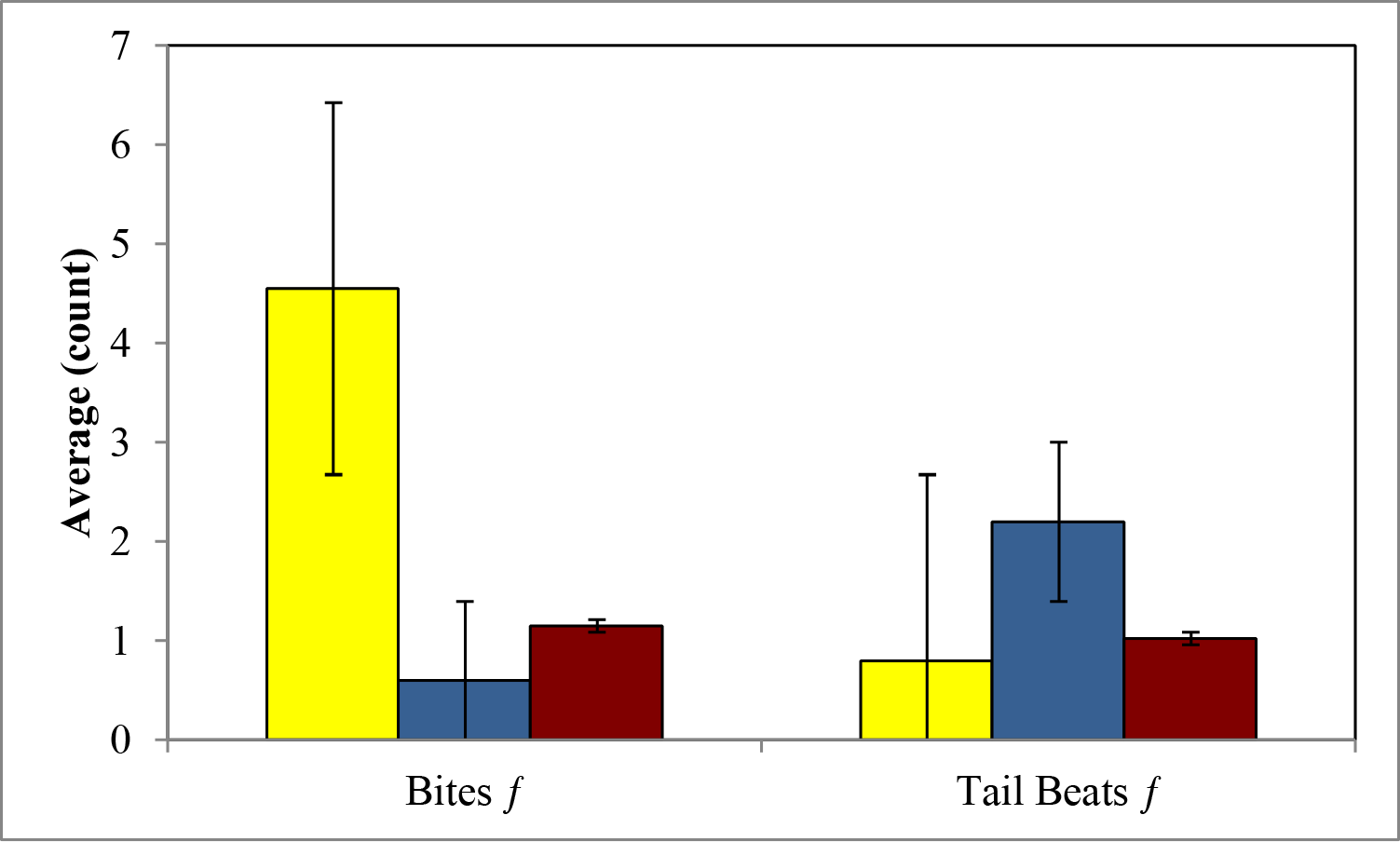
Average±SE amount of aggression in bites and tail beatsthat focal females received from partner females: heterospecific females (yellow), clonal sisters (blue), and non-sister clones (red). Focal femalesreceived more bites from heterospecific females and received more tail beats from clonal sisters.

**Fig. 3.**
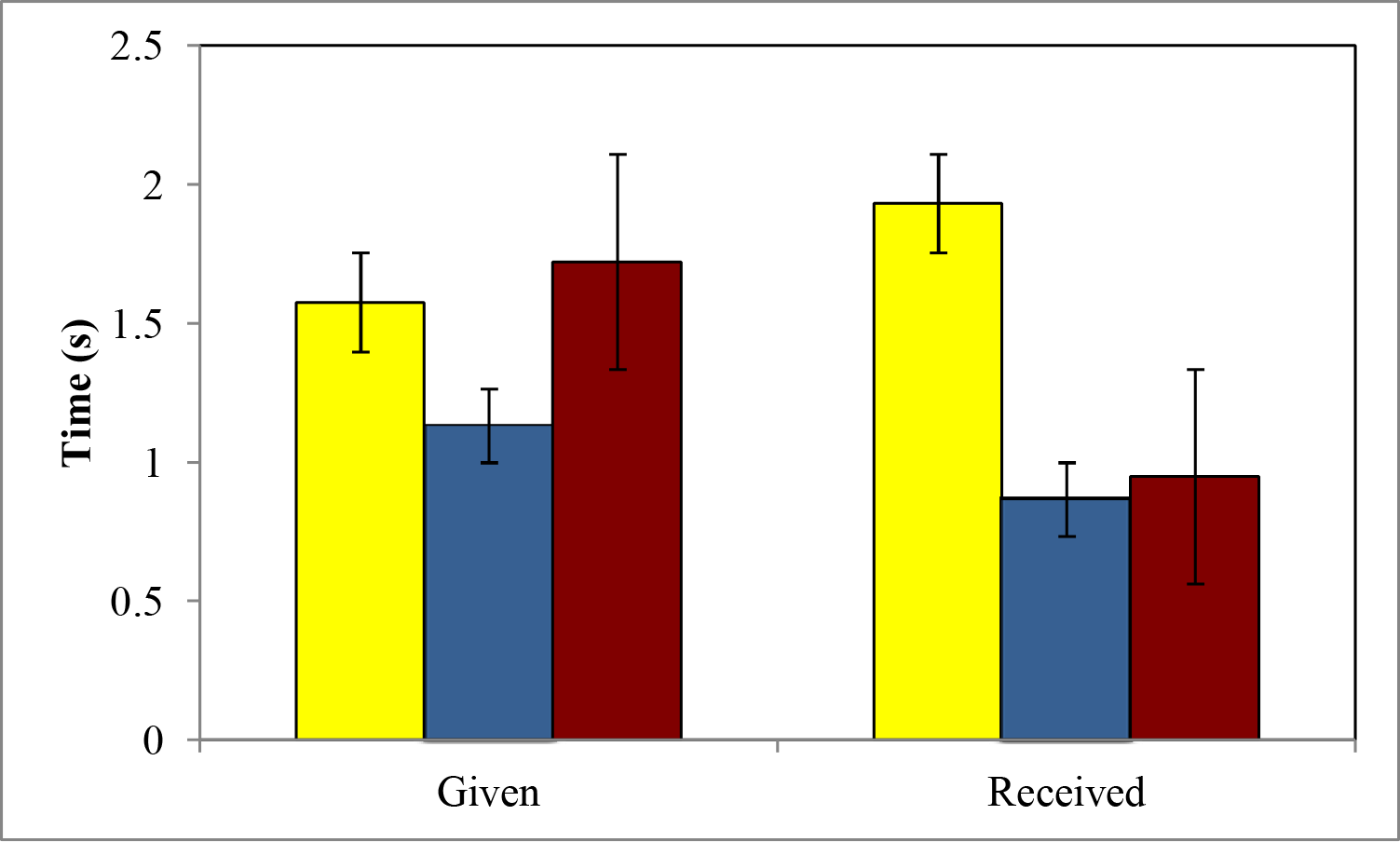
Average time (s)±SE of aggression focal females gave and received from partner females: heterospecific females (yellow), clonal sisters (blue), and non-sister clones (red). The average time a focal female spent behaving aggressively was higher when paired with either a heterospecific female or a non-sister clone when compared to clonal sisters. The average amount a focal female received aggression was higher when paired with aheterospecific than either a clonal sister or non-sister.

**Table 3:** Average±SD of aggressive behaviors both given and received 465 in the three treatments.

| Treatment | Aggression Given |  |  | Aggression Received |  |  |
| --- | --- | --- | --- | --- | --- | --- |
|  | Time (s) | Bites | Tail beats | Time (s) | Bites | Tail beats |
| Heterospecific Partner | 1.653 ± 3.17 | 2.11 ± 4.51 | 2.82 ± 6.05 | 1.936 ± 3.59 | 4.53 ± 8.48 | 0.84 ± 1.31 |
| Conspecific Clonal Sister Partner | 1.2 ± 2.49 | 2.05 ± 2.84 | 1.08 ± 2.16 | 0.9 ± 1.72 | 0.63 ± 1.36 | 2.32 ± 4.97 |
| Conspecific Non-Sister Clone Partner | 1.737 ± 3.79 | 3.84 ± 6.51 | 1.63 ± 3.45 | 0.972 ± 2.97 | 1.16 ± 3.01 | 0.87 ± 1.56 |

## Discussion

In the present study, we evaluated female-female aggression in Amazon mollies towards both heterospecific and conspecific females, and clonal sisters and non-sister clones. We were able to confirm that Amazon mollies do, indeed, adjust their aggressive behaviors towards other females in their social environment (Makowicz and Schlupp 2015, Makowicz et al. 2016). In addition, we were able to show that in a food-limited environment, females maintained this ability to adjust their aggressive behaviors relative to relatedness.

The aggression in Amazon mollies was not as intense as when females were sated (Makowicz et al. 2016) or as intense as in other fish species (Arnott and Elwood 2009a; Brown and Brown 1996; Olesen and Jarvi 1997). Nonetheless, focal females spent more time behaving aggressively to both heterospecific females and non-sister clones, however, they only gavemore bites to non-sister clones. Because focal females were receiving significantly more aggressive bites from the heterospecific female, this may have resulted in the reduction we see in the focal females returning the behavior; although the focal female did perform more tail beats towards the heterospecific female. Indeed, Amazon mollies received less aggression from either conspecific female when compared to the heterospecific sailfin molly female. Although, we did not take into consideration prior dominance interactions that may have occurred prior to isolating the individuals, which may influence female aggression (Laskowski et al. 2016), randomly selecting and isolating individuals from all threestock populations would havecontrolled for any prior dominance effects that we found in our fish.

Unlike the time females spent behaving aggressively and the number of bites females gave, females received more tail beats from clonal sisters than either non-sister clones or heterospecific females. It is possible that tail beats may act as a pre-aggressive behavior in this species and serve as a possible “warning” signal rather than an act of full aggression. Unlike bites where females can remove scales or chasing where there is a high metabolic cost to flee, tail beats may not be as costly to an opponent. While tail beats in some species may be rather intense (*Nannacara anomala;* Hurd 1997) and are more intense when performed by males when compared to females (Arnott and Elwood 2009a; LaManna and Eason 2010), in general, tail beats are typically a low risk behavior when compared to head-on aggressive behaviors (Arnott and Elwood 2009b; Ros et al. 2006). Also, the metabolic demand for tail beats is not as costly as other aggressive behaviors, such as the overt aggression and lateral displays performed by males (Ros et al. 2006). Indeed, together, this suggests that tail beats are a low cost, warning signal, rather than an intense aggressive behavior, which was previously thought for this species.

Similar to other kinship studies, Amazon mollies adjusted their aggressive behaviors towards related females when resources are limited. Females spent less time performing aggressive behaviors to clonal sisters when compared to non-sisters. Interestingly, there was no difference in the numberof bites Amazon females performed towards clonal sisters and nonsisters, although they still received more aggressive bites from non-sister clones when compared to clonal sisters. This seems to be a result of the induced hunger of the focal females. When Amazon females were sated prior to aggressive trials, focal females exhibited little to no aggression towards, nor did they receive any aggression from, clonal sisters (Makowicz et al. 2016). Thus, hunger seems to reduce the discrimination of females when performing bites, but not the amount of time females spent behaving aggressively or performing tail beats.

This study suggests that Amazon mollies are indeed more altruistic and less antagonistic to clonal sisters when compared to non-sisters by adjusting their aggressive behaviors when in a low food environment. While they performed an equivalent number of bites to both clonal sisters and non-sister clones, only the overall time spent being aggressive was affected by hunger. Nonetheless, other social environment conditions such as audience effects (either heterospecific or conspecific, male or female, or clonal sister or non-sister), the presence of predators, the reproductive state of a female, or the availability of refuges may influence the intensity of female-female aggression in this species.

## Acknowledgements

We would like to thank Shelby Burridge and Elizabeth Hardy for their input on previous versions of this manuscript. This research was approved by the Institutional Animal Care and Use Committee (R13-006). The Texas Parks and Wildlife department kindly issued a research and collection permit # SPR-0305-045.

